# Automated Segmentation of Hepatic Vessels and Lobules in Whole-Slide Images Using U-Net Models

**DOI:** 10.1101/2025.09.08.674181

**Authors:** Mehul Bafna, Matthias König, Sylvia Saalfeld, Vladimira Moulisova, Vaclav Liska, Uta Dahmen, Mohamed Albadry

## Abstract

Automated analysis of hepatic vascular structures and lobules within whole-slide histological images is critical for ensuring accurate and timely morphometric evaluations and facilitating advancements in computational liver histology. Nonetheless, the intricate morphology of the tissue, variability in staining techniques, and the requirements for high-resolution imaging present substantial challenges to the precision of segmentation processes. We present a robust deep-learning pipeline using adaptive patch extraction and specialized U-Net architectures for segmenting vessels, bile ducts, and lobules in Glutamine Synthetase and Picro-Sirius-Red stained porcine liver sections. Our architecture incorporates a weight-boosted nnU-Net framework to effectively manage class imbalances and improve the detection of smaller vascular structures. Geometric data transformations enhanced the robustness and generalizability of the segmentation models. Evaluations conducted through five-fold cross-validation, as well as assessments utilizing independent test datasets, resulted in Dice similarity scores: 0.960 for lobules, 0.801 for central veins, 0.909 for hepatic arteries, 0.609 for portal veins, and 0.710 for bile ducts. The developed segmentation pipeline additionally supports comprehensive morphometric analyses of structural parameters, including number and size (diameter, area) of vascular structures, bile ducts, and lobules. For e.g., the diameter of hepatic arteries ranges between 30-90 *µ*m. These findings underscore the practical relevance of adaptable segmentation frameworks in advancing computational histological analysis of liver tissue.

## 1 Introduction

The liver is a highly vascularized organ responsible for critical physiological functions, including metabolism, detoxification, and immune regulation [ADEK19]. Key histological features of the liver include vascular components—such as portal veins, central veins, and hepatic arteries—and non-vascular structures like bile ducts, all of which are organized within hepatic lobules, the fundamental structural and functional units of the liver [TGW17]. Prior work has examined structural elements such as hepatic lobuli, without focusing on vascular structures [AKG^+^24].

Advances in digital histology and whole-slide imaging (WSI) have enabled automated analysis of liver tissue. However, accurately segmenting both vascular structures and lobular boundaries simultaneously remains challenging. Liver tissue segmentation presents several challenges due to the complex anatomical and histological characteristics of hepatic structures. Low contrast regions, often caused by inconsistent staining and focal tissue artifacts, can obscure the boundaries between vessels and surrounding parenchyma, making the delineation of small arteries and bile ducts particularly difficult [CS16]. Additionally, the wide range of scales among vascular structures introduces further complexity, as a segmentation approach must capture the fine-grained details of small vessels while also accurately identifying larger, branching structures. Performance often varies with size: smaller vessels, being contained within individual image patches, are typically segmented more accurately, whereas larger vessels that span multiple patches are susceptible to fragmentation and misclassification. The morphological similarity between different hepatic components, such as vascular structures and bile ducts, further increases class ambiguity, complicating accurate differentiation and lobular recognition [Inv13]. Moreover, effective segmentation is heavily dependent on contextual cues, as structures like portal fields and central veins provide critical spatial references. Due to these difficulties, most of the previous studies focus either on lobule detection or the identification of vascular structures, which limits their applicability.

Most existing vascular segmentation studies targeting the liver focus on radiological modalities, particularly Magnetic Resonance Imaging (MRI) and Computed Tomography (CT). A thorough examination of various computational methods for liver vessel segmentation, encompassing both traditional image processing and modern deep learning techniques, is provided in the review [CK21]. The effectiveness of nnU-Net and its variants for liver segmentation tasks has been demonstrated prior in MRI and CT images [HJH^+^24, HAT^+^23]. Nevertheless, the research gap remains limited for vascular segmentation within histological images.

Therefore, we propose a regularized U-Net architecture to segment both hepatic vascular and non-vascular structures as well as lobular boundaries effectively in whole slide images from histological liver sections. Integrating deep learning into the segmentation framework enhances the ability to delineate intricate vascular and non-vascular structures and heterogeneous lobular patterns, thereby improving accuracy across multiple spatial scales.

The porcine liver utilized in the study serves as an ideal experimental model due to the clear delineation of hepatic lobules by collagen-rich septae, facilitating accurate segmentation of vascular structures and offering essential contextual information for defining lobular organization [MSN^+^12]. It provides an effective foundation for the future development of segmentation algorithms that are transferable across species and adaptable to diverse histological conditions.

## 2 Materials and Methods

### 2.1 Dataset Preparation

Liver samples were explanted from Prestice black-pied pigs (weighing 25–33 kg) under general anesthesia and immediately fixed in formalin with no ischemia time. The work with animals was conducted under the law of the Czech Republic, which is in line with the legislation of the European Union. The procedure protocol was approved by the Ministry of Education, Youth and Sport of the Czech Republic (approval no. MSMT-15629/2020-4).

A single whole slide image (WSI) of a liver biopsy from a clinically healthy, 3-month-old female Prestice Black-Pied pig was selected for model development. Four additional WSIs, obtained from different clinically healthy Prestice Black-Pied pigs (both male and female) of the same age and weight range, were used for testing. All tissue sections were obtained from archived formalin-fixed and paraffin-embedded liver tissue blocks.

Briefly, liver sections were cut at 3*µ*m thickness and processed using a standardized immunohistochemistry protocol. Immunostaining was performed using a mouse monoclonal anti-glutamine synthetase (GS) antibody (MAB302; 1:10,000 dilution; Merck, Germany), followed by counterstaining with Picro-Sirius Red (PSR; 1342500250, Morphisto, Germany). Detection was carried out using an avidin-biotin system consisting of a biotinylated goat anti-mouse IgG H&L (ab6788, Abcam, Germany), Streptavidin-HRP (ab64269, Abcam, Germany), and an avidin/biotin blocking kit (ab64212, Abcam, Germany). DAB chromogen (GV825, Dako, Denmark) was used for signal visualization.

GS staining reveals a characteristic perivenous hepatocyte distribution in the liver, aiding in hepatic zonation analysis [Nor79]. PSR staining was performed as counterstain which produces a distinctive red coloration of collagen fibers under brightfield microscopy [LYL^+^14]. In addition, it provides excellent contrast for vessel wall delineation compared to standard HE staining [JBB79].

Slides were digitized at a resolution: 227 nm/pixel using a Hamamatsu NanoZoomer scanner (model L11600) with NDP.view2 software. The scanned WSI used in the data modeling had dimensions of 111,104 *×* 67,072 pixels. Annotations were created using the Hamamatsu NDP.view2 annotation toolbox by a trained lab technician irrespectively of size. Due to the large WSI size, training was conducted on smaller patches. The WSI was divided into non-overlapping patches using the OpenSlide module (version 1.2.0) in Python to facilitate training [GGH^+^22].

#### 2.1.1 Vessel Segmentation Dataset

For vessel segmentation, image patches were extracted at a size of 992 × 1048 pixels and then padded to 1056 × 1056 pixels to match the input dimension requirements of the nnU-Net architecture. This padding ensured compatibility with the downsampling and upsampling operations (based on a factor of 2^*n*^, where n=5) of the network and helped prevent information loss at patch boundaries [IPK^+^18]. All patches were saved in Tagged Image File Format (TIFF) with lossless compression to preserve image quality.

The vessel segmentation dataset consisted of 3,574 non-overlapping patches extracted from a single WSI. A random 10% (357 patches) was reserved for testing, while the remaining 3,217 patches (90%) were used for five-fold cross-validation. In each fold, 2,574 patches were used for training and 643 for validation with random horizontal/vertical flips and rotations (*±*30^*◦*^) applied for data transformation. Each image patch was paired with a manually annotated segmentation mask containing five classes: background (black, pixel value 0), portal vein (blue, value 1), central vein (green, value 2), hepatic artery (red, value 3), and bile duct (purple, value 4) (refer to **Figure 1**). The masks were saved in TIFF format with the same spatial dimensions and padding as the corresponding image patches.

**Figure 1.**
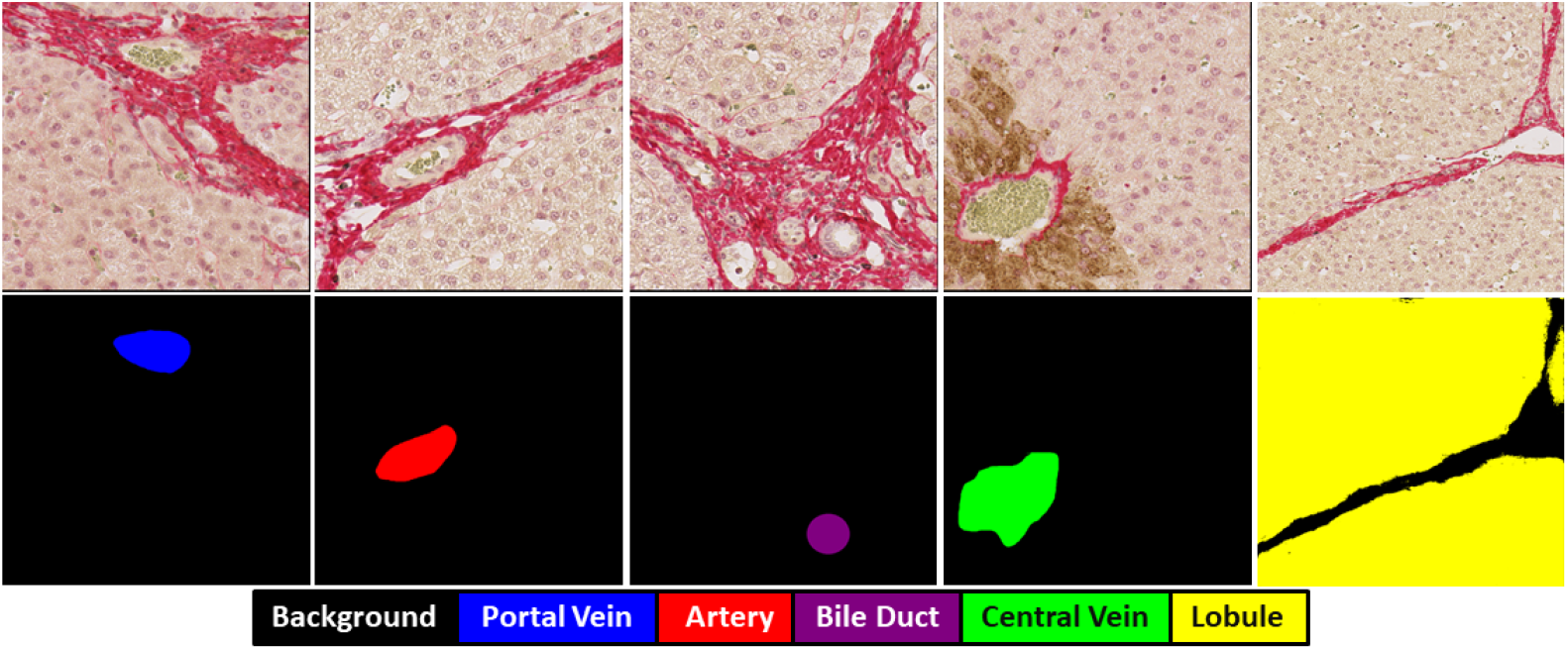
Overview of original image and ground truth segmentation. From top to bottom: Original patch and its corresponding ground truth multi-class mask.

#### 2.1.2 Lobule Detection Dataset

For hepatic lobule detection, a separate dataset of 1055 non-overlapping patches was prepared with a larger dimension of 1760 *×* 2112 pixels. The increased patch size was specifically chosen to encapsulate entire hepatic lobules within single images, as lobules represent larger structural units compared to individual vessels. Each lobule patch was saved in TIFF format with corresponding binary masks, where yellow represents lobule regions as shown in **Figure 1**. Similar to the hepatic vascular segmentation model, a random 10% split was used to select 105 patches for testing. The remaining 950 patches (90% of the dataset) were used for five-fold cross-validation. In each iteration, 760 patches (four folds) were used for training and 190 patches (one fold) for validation, applying the same geometric data transformations as in the vessel segmentation model.

The complete end-to-end dataset preparation pipeline is illustrated in **Figure 2**. The class distribution in the dataset for Network 1 was highly imbalanced, with background occupying 96.8% of the total pixels (refer to **Table 1**). This strong class imbalance was decisive for the choice of loss function and data augmentation strategies.

**Table 1:** Overview of annotated structures and their pixel-wise representation in vessel and lobule segmentation networks.

| Structure type | Total structures annotated | Pixel proportion<br>(% of total image pixels) |
| --- | --- | --- |
| **Network 1: Vessel Segmentation** |  |  |
| Background | - | 96.8% |
| Portal vein | 714 | 2.1% |
| Artery | 229 | 0.35% |
| Bile duct | 350 | 0.25% |
| Central vein | 173 | 0.5% |
| **Network 2: Lobule Detection** |  |  |
| Background | - | 65.3% |
| Lobule | 225 | 34.7% |

**Figure 2.**
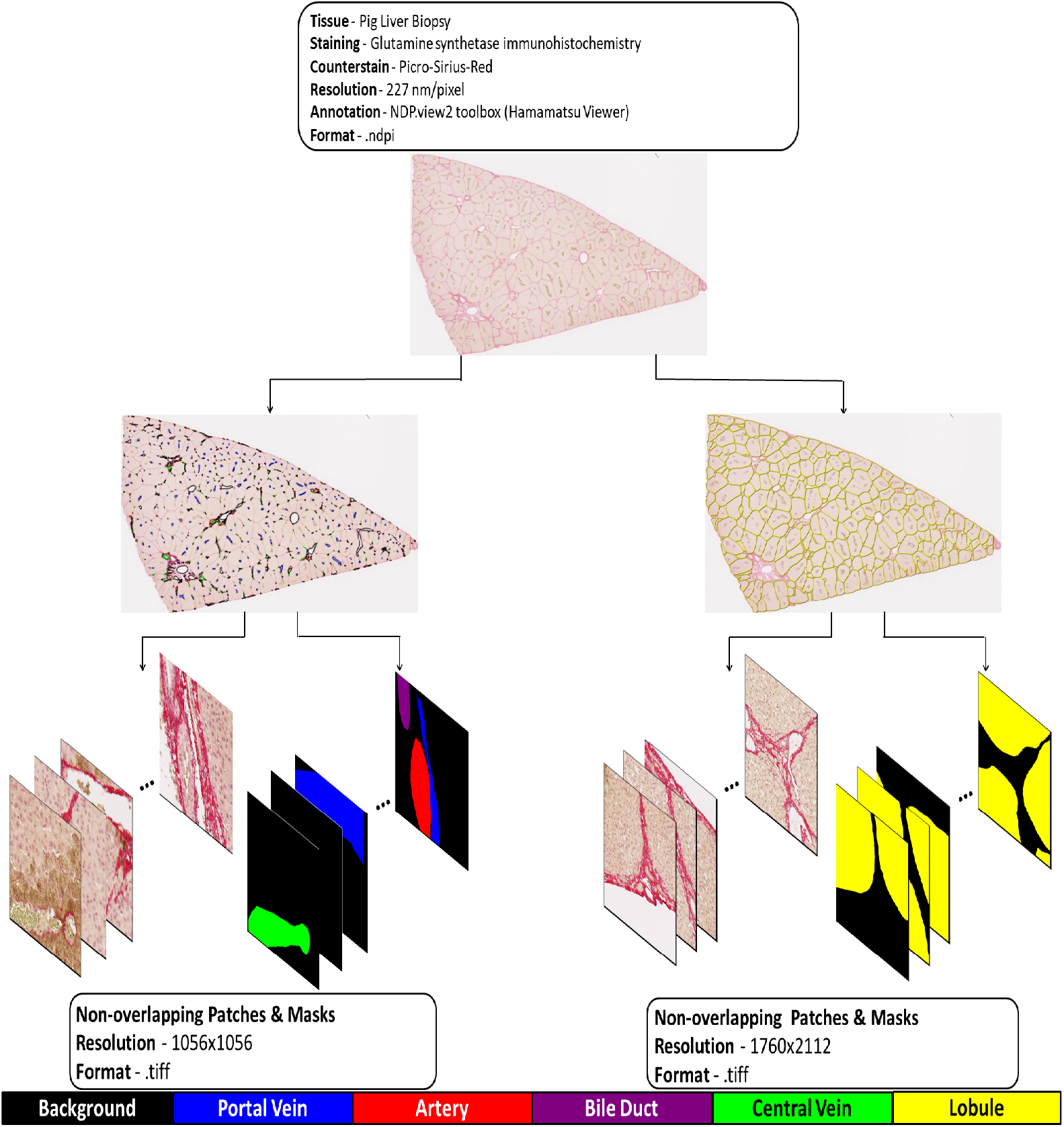
Stepwise breakdown of dataset preparation. Dataset Preparation Workflow: Stage 1 – WSI Generation; Stage 2 – Separate Annotations for Hepatic Vessel Segmentation Training and Lobule Segmentation Training; Stage 3 – Extraction of Annotated Patches and Corresponding Ground Truth Masks from WSIs.

### 2.2 Network Architecture

We propose two state-of-the-art, distinct segmentation models: a weight-boosted nnU-Net architecture for liver vessel segmentation and a standard nnU-Net model for hepatic lobule detection. Separate networks for vessel and lobule segmentation were employed to prevent feature confusion and improve specificity, particularly since central veins are located within the hepatic lobule.

#### 2.2.1 Vessel Segmentation Network

The vessel segmentation network is based on the nnU-Net architecture [IPK^+^18] integrated with an adaptive weight-boosting mechanism to address the strong class imbalance between vessel and background pixels. The model processes input images of size 1056×1056×3 through an encoder with four blocks, each containing a double convolutional layer (DoubleConv) followed by max-pooling, progressively increasing feature channels (32*→*64*→*128*→*256) and reducing spatial dimensions (refer to **Figure 3**). The encoder ends with a bottleneck layer (66×66×512) for feature processing. The decoder mirrors the encoder, with upsampling blocks that sequentially reduce feature channels (256*→*128*→*64*→*32) while restoring spatial details. The weight-boosting mechanism dynamically adjusts the class-wise loss contributions during training, ensuring proper learning despite the underrepresented vessel class.

**Figure 3.**
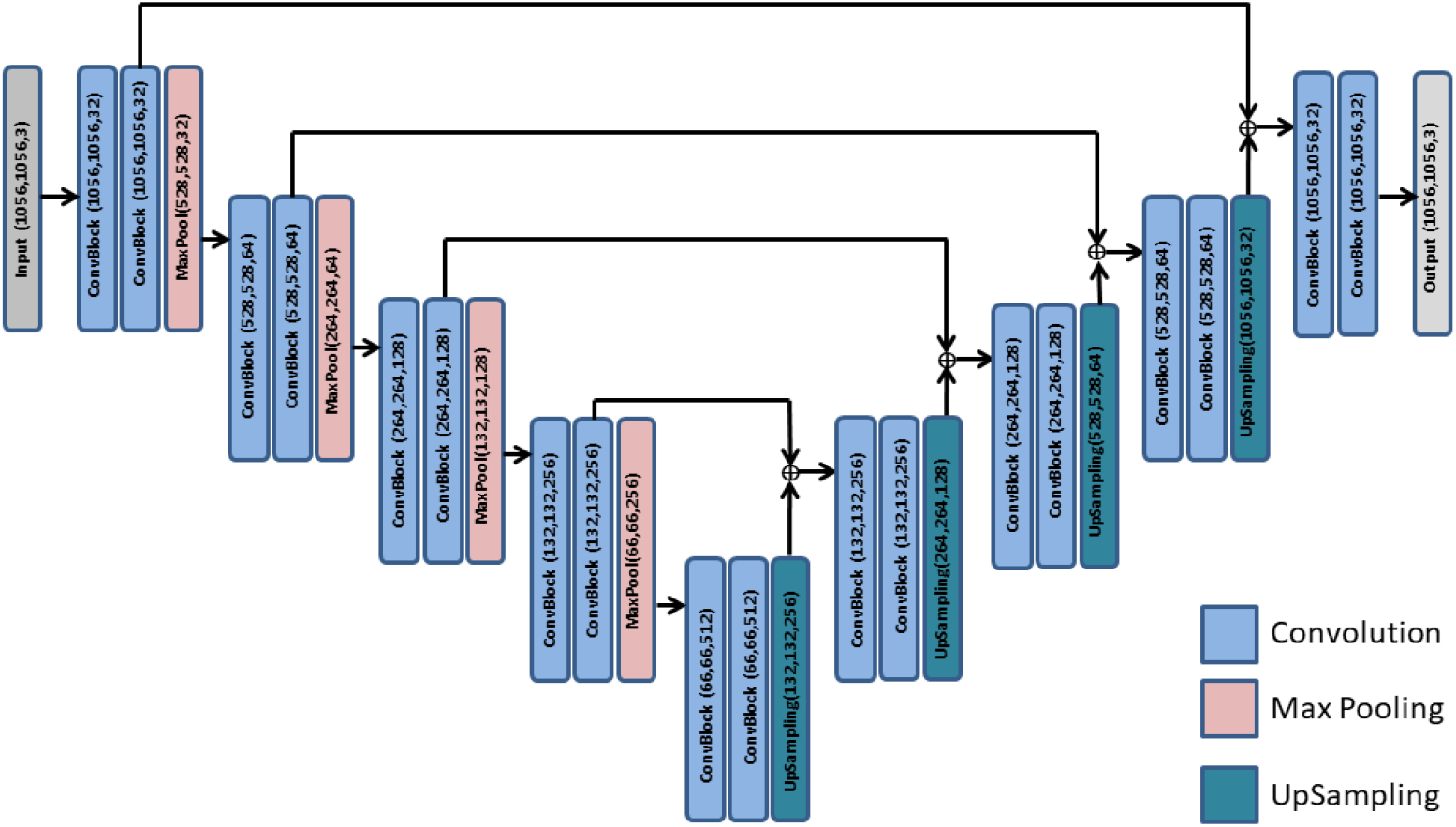
Model architecture used for segmentation tasks. Architecture of the nnU-Net model used in vessel segmentation and lobule detection.

#### 2.2.2 Lobule Detection Network

The second network focuses on hepatic lobule detection and is built upon the same nnU-Net architectural framework. Unlike the vessel segmentation network, this model does not implement weight-boosting mechanisms as the class distribution is less severely imbalanced. The lobule detection network maintains the same encoder-decoder structure with four blocks in each path and identical feature channel progression. Skip connections between corresponding encoder and decoder layers facilitate the preservation of spatial information throughout the network, which is essential for accurately delineating lobular boundaries.

Although both networks leverage the nnU-Net framework, the key difference lies in the optimization strategy, with the vessel segmentation network specifically adjusted to handle the challenges of segmenting fine vascular structures that constitute a small fraction of the overall image area.

### 2.3 Experiments

The experiments were conducted on the high-performance computing (HPC) cluster in Jena, Germany, equipped with NVIDIA A100 graphical processing units (GPUs). The software environment comprised Python version 3.10, PyTorch version 2.6, and CUDA 11.8.

Both U-Net variants were trained with the following hyperparameters:

- **Epochs**: 250
- **Batch size**: 8
- **Optimizer**: Adam’s optimizer with weight decay 0.0002
- **Learning rate**: 0.0005 (Cosine Annealing)

Cosine annealing is a learning rate scheduling strategy that smoothly decreases the learning rate following a cosine curve. This gradual decay helps the model converge more effectively by avoiding abrupt changes in the learning rate. The learning rate *η*_*t*_ at epoch *t* is given by the formula:

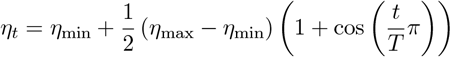

where:

- *η*_max_ is the initial (maximum) learning rate,
- *η*_min_ is the minimum learning rate,
- *T* is the total number of epochs or iterations.

To address the class imbalance inherent in the whole-slide images where background pixels substantially outnumber vessel pixels, an adaptive weight-boosting mechanism was applied during the training of the hepatic vessel segmentation model. The algorithm dynamically adjusted class weights every 15 epochs based on class-wise Dice scores: for classes with Dice scores below 0.3, weights were multiplied by a factor of 1.3; for classes with scores between 0.3 and 0.5, weights were multiplied by 1.2; and for classes with scores between 0.5 and 0.7, weights were multiplied by 1.1. For vessel segmentation, a weighted combination of Dice, focal, and cross-entropy losses was applied. In contrast, the lobule detection model utilized a simpler binary cross-entropy with Dice loss formulation, as the pixel distribution was less imbalanced (refer to **Figure 4**).

**Figure 4.**
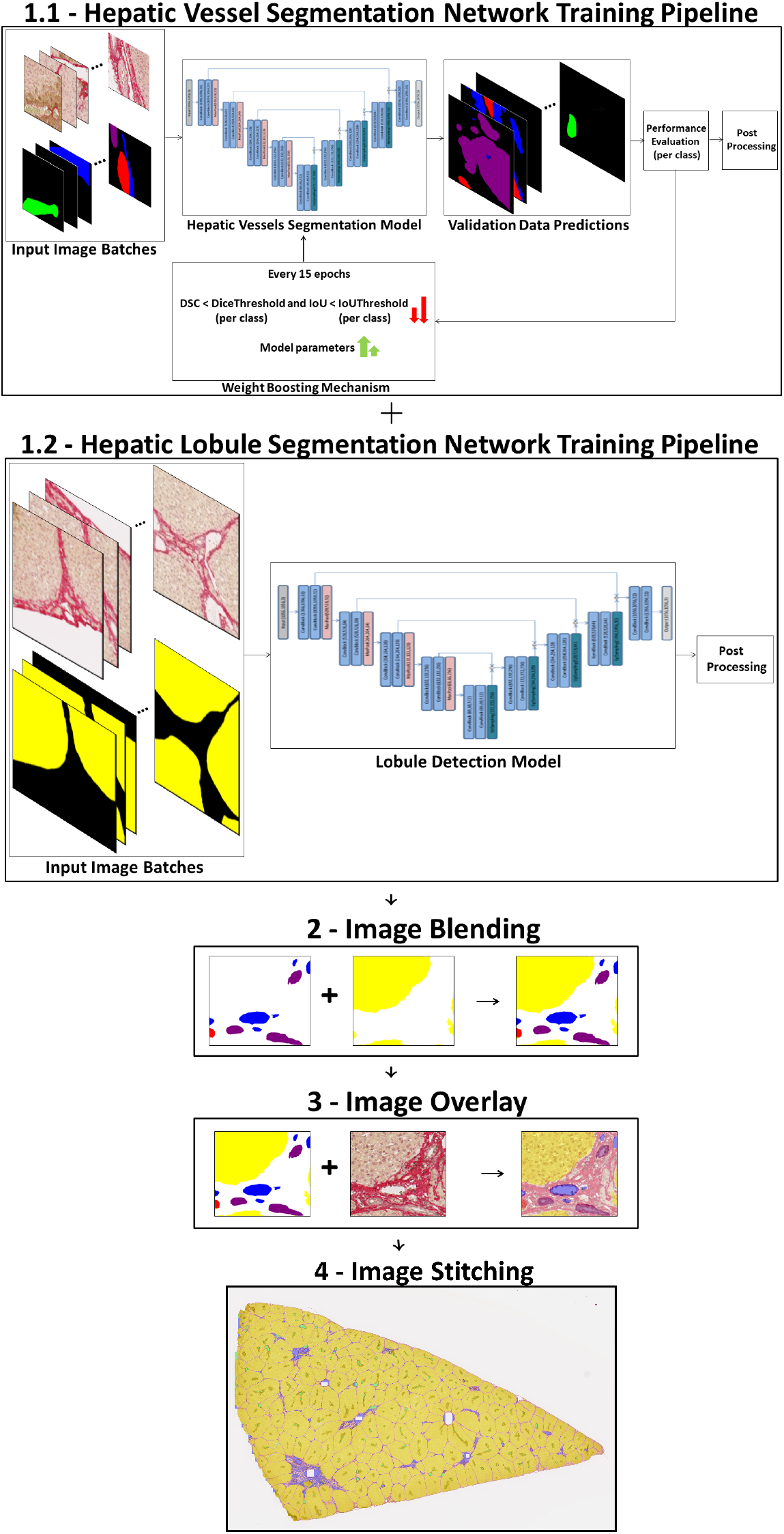
End-to-end pipeline for segmentation and integration. Overview of the end-to-end dataset preparation pipeline for hepatic tissue analysis.

Additionally, to handle overfitting, an early stopping mechanism was implemented, terminating training if the validation loss showed no improvement over 20 consecutive epochs. Furthermore, 5-fold cross-validation was employed to ensure robust evaluation and generate reliable final results across the complete dataset.

### 2.4 Metrics

To evaluate the segmentation performance, we utilized the Dice Similarity Coefficient (Dice score), Intersection over Union (IoU), True Positive Rate (TPR), and Average Symmetric Surface Distance (ASSD). These metrics are widely used in medical image analysis to quantify overlap and accuracy between predicted and ground truth segmentations.

#### 2.4.1 Dice Similarity Coefficient (DSC)

The Dice score measures the overlap between the predicted segmentation P and the ground truth G, and is defined as:

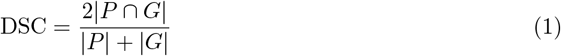

For a specific class c, the class-wise Dice score is given by:

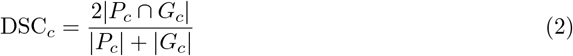

#### 2.4.2 Intersection over Union (IoU)

IoU, also known as the Jaccard index, is defined as the ratio of the intersection over the union of the predicted and ground truth regions:

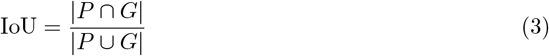

For a specific class c, the class-wise IoU is given by:

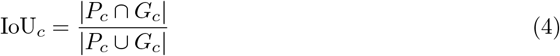

#### 2.4.3 True Positive Rate (TPR)

The True Positive Rate measures the proportion of actual pixels for a specific class that are correctly identified by the model. Given a predicted segmentation *P* and the ground truth *G*, the TPR is defined as:

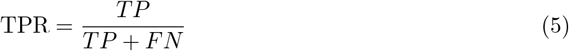

For a specific class c, the class-wise TPR is given by:

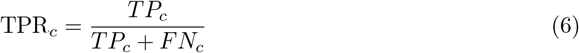

where *TP* denotes the number of true positives (correctly predicted positive pixels) and *FN* is the number of false negatives (missed positive pixels) for that class.

#### 2.4.4 Average Symmetric Surface Distance (ASSD)

The Average Symmetric Surface Distance quantifies the mean distance between the surface boundaries of the predicted segmentation P and the ground truth G. It is computed as the average of distances from each point on one surface to the closest point on the other surface, considering both directions to ensure symmetry:

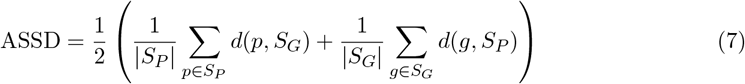

where *S*_*P*_ and *S*_*G*_ represent the surface boundaries of the predicted and ground truth segmentations respectively, |*S*_*P*_ | and |*S*_*G*_| denote the number of surface points, and *d*(*p, S*_*G*_) represents the minimum Euclidean distance from point *p* to any point on surface *S*_*G*_.

For a specific class c, the class-wise ASSD is given by:

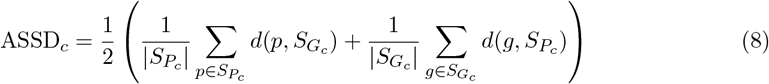

The Dice score and IoU provide insights into the degree of region overlap, while TPR highlights the completeness of the segmentation by measuring how well positive regions are detected. ASSD complements these surface metrics by quantifying boundary precision and evaluating the spatial accuracy of lobule contours.

### 2.5 Morphometric Analysis

The overall stitched images compiled either from vessel or from lobule segmentation models were postprocessed prior to morphometric analysis. Determination of the surface area of all vessel structures did not require any further morphometric operations. In contrast, the complete lobule image underwent two morphological operations to avoid underestimation of lobule number due to incomplete separation by septae. Dilation was used to widen the septae and erosion to close eventual remaining gaps. While dilation effectively expanded the septal boundaries, the subsequent erosion process, designed to close remaining discontinuities, resulted in a net contraction of the overall lobular area. Surface areas of the resulting objects representing the lobules was calculated by counting the number of pixel for each object.

On both images, area size filters with ranges were applied to reduce the number of false positive counts. For lobules, the size range was set from 12500*µm*^2^ - 1250000*µm*^2^. The upper range was selected to exclude falsely connected lobules. For portal veins the size range was set to at least 1000*µm*^2^, for hepatic artery and bile duct, at least 125*µm*^2^ and for central vein, at least 125*µm*^2^. In order to not exclude the large vessels we did not set an upper range. After excluding eventually falsely detected small or very large objects such as interconnected lobules, the total number of all structures within the given area size was determined, for lobules using the lobule segmentation model and vessel structures using the vessel segmentation model. To better quantify the shape of the accepted structures, we computed the perimeter by considering the covered area as an equivalent circle. Furthermore, we determined both Feret maximum and minimum diameter. Feret or caliper diameter refers to the distance between two parallel lines tangent to the projected contour.

#### Major Feret Diameter

The Feret diameter major axis represents the maximum distance between any two points on the boundary of a structure when measured in all possible directions. This measurement captures the longest dimension of the object, providing insight into its overall size and orientation.

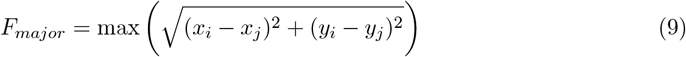

where (*x*_*i*_, *y*_*i*_) and (*x*_*j*_, *y*_*j*_) are all possible pairs of boundary points.

#### Minor Feret Diameter

The Feret diameter minor axis represents the minimum distance between parallel lines that can be drawn to just touch opposite sides of the structure. This measurement captures the narrowest dimension of the object, complementing the major axis to provide a complete picture of the dimensional characteristics for the given structure.

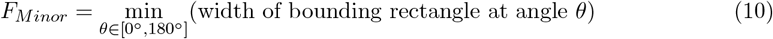

where *θ* ranges from 0° to 180°, and the width is measured perpendicular to the longest side at each angle. An illustration of the feret diameters for a segmented central vein is presented in **Figure 5**.

**Figure 5.**
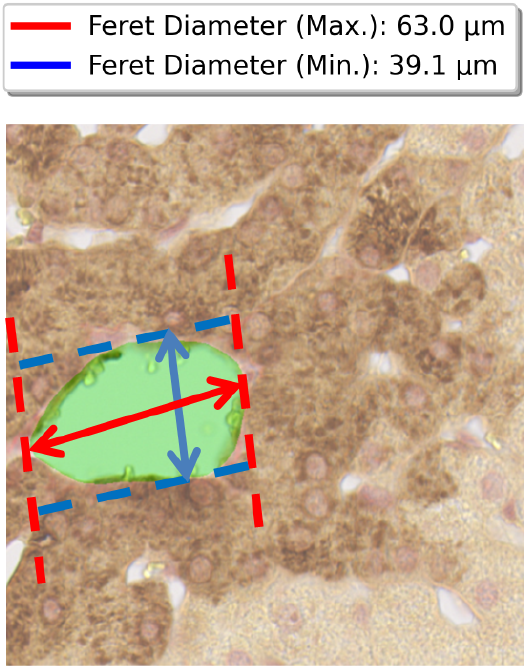
Feret diameter measurements for central vein. Red arrow indicates major axis (maximum boundary distance), blue arrow indicates minor axis (minimum width between parallel lines). Measurements converted to micrometers using 0.227 pixel scaling factor.

Aspect ratio was calculated by dividing minimum and maximum Feret diameter.

### 2.6 Code and Data Availability

To support reproducibility for further research, we have made all relevant resources publicly available. The complete implementation—including deep learning models, data preprocessing scripts, and evaluation tools is hosted on our GitHub repository: https://github.com/MehulBafna/Image-Segmentation-Paper. The repository is organized to reflect each stage of the pipeline and is released under an open-source license.

Our annotated dataset, which includes whole slide images, corresponding image patches, and multiclass segmentation masks for hepatic vessels and lobules, is available through Zenodo: https://doi.org/10.5281/zenodo.17041227.

## 3 Results

### 3.1 Performance Metrics

Model performance was evaluated using 5-fold cross-validation (see **Table 2**) as well as results from the test data (see **Table 3**). The results presented in **Table 2** indicate the mean over all the image patches per fold and subsequently over all five cross-validation folds. The nnU-Net achieved mean dice scores in the range 0.74 - 0.84 for hepatic vessel segmentation across portal vein, central vein, artery, and bile duct structures. For lobule segmentation, the model achieved a higher mean Dice score in the range of 0.96 - 0.97. Detailed cross-validation metrics are summarized in **Table 2**.

**Table 2:**
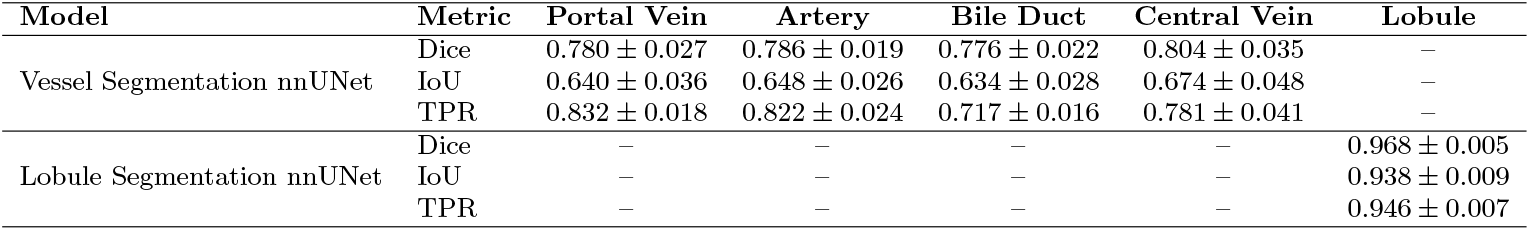
Performance Metrics (Dice vs IoU vs TPR) with 5-fold cross validation.

**Table 3:**
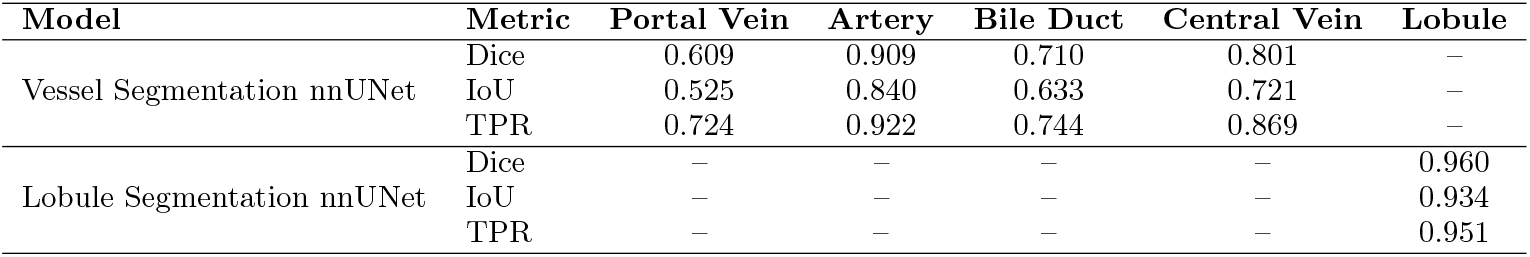
Performance Metrics (Dice vs IoU vs TPR) on the test data.

| Model | Metric | Portal Vein | Artery | Bile Duct | Central Vein | Lobule |
| --- | --- | --- | --- | --- | --- | --- |
| Vessel Segmentation nnUNet | Dice | 0.609 | 0.909 | 0.710 | 0.801 | – |
|  | IoU | 0.525 | 0.840 | 0.633 | 0.721 | – |
|  | TPR | 0.724 | 0.922 | 0.744 | 0.869 | – |
| Lobule Segmentation nnUNet | Dice | – | – | – | – | 0.960 |
|  | IoU | – | – | – | – | 0.934 |
|  | TPR | – | – | – | – | 0.951 |

On the test dataset, the nnU-Net achieved mean dice scores in the range of 0.61 - 0.91 across four vascular structures—portal veins, central veins, hepatic arteries, and bile ducts. Furthermore, lobule segmentation attained a higher mean Dice score of 0.960. The detailed results are provided in **Table 3**, and exemplary segmentations are shown in **Figure 7**.

**Figure 6.**
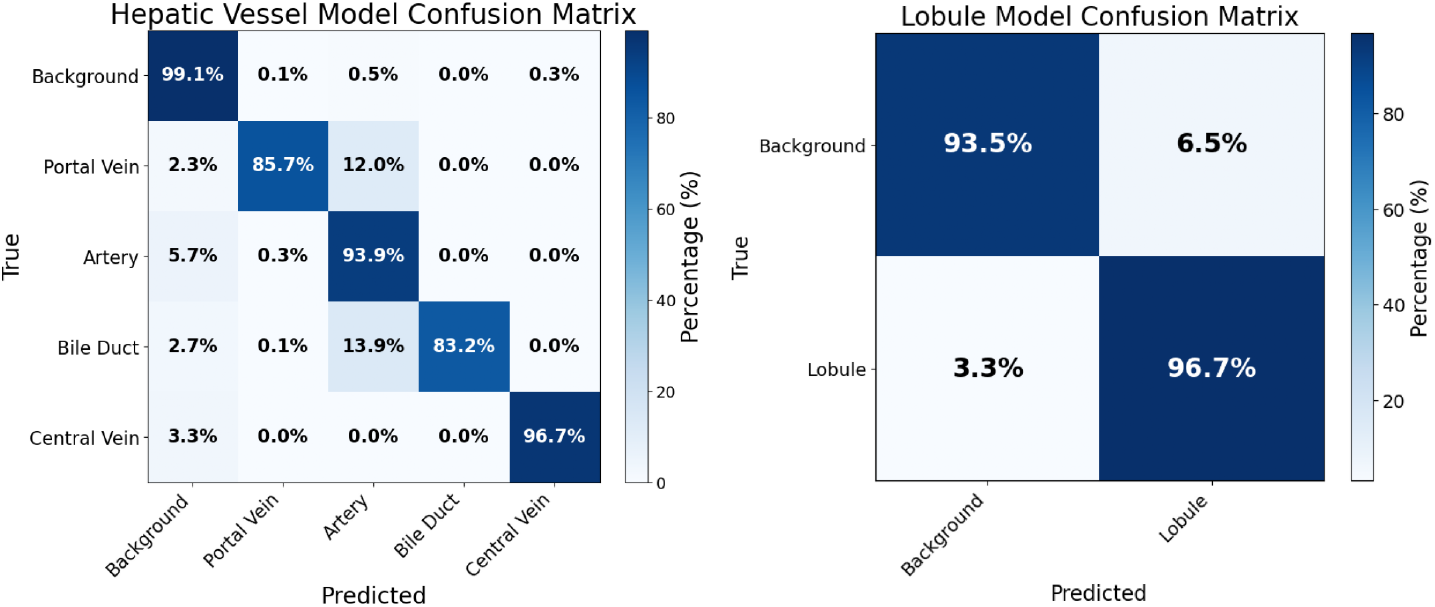
Confusion matrices for hepatic vessel and lobule segmentation models. The hepatic vessel model (left) shows classification performance across five classes: Background, Portal Vein, Artery, Bile Duct, and Central Vein. The lobule model (right) demonstrates binary classification performance between Background and Lobule structures. Values represent percentages of true class predictions, with darker blue indicating higher accuracy.

**Figure 7.**
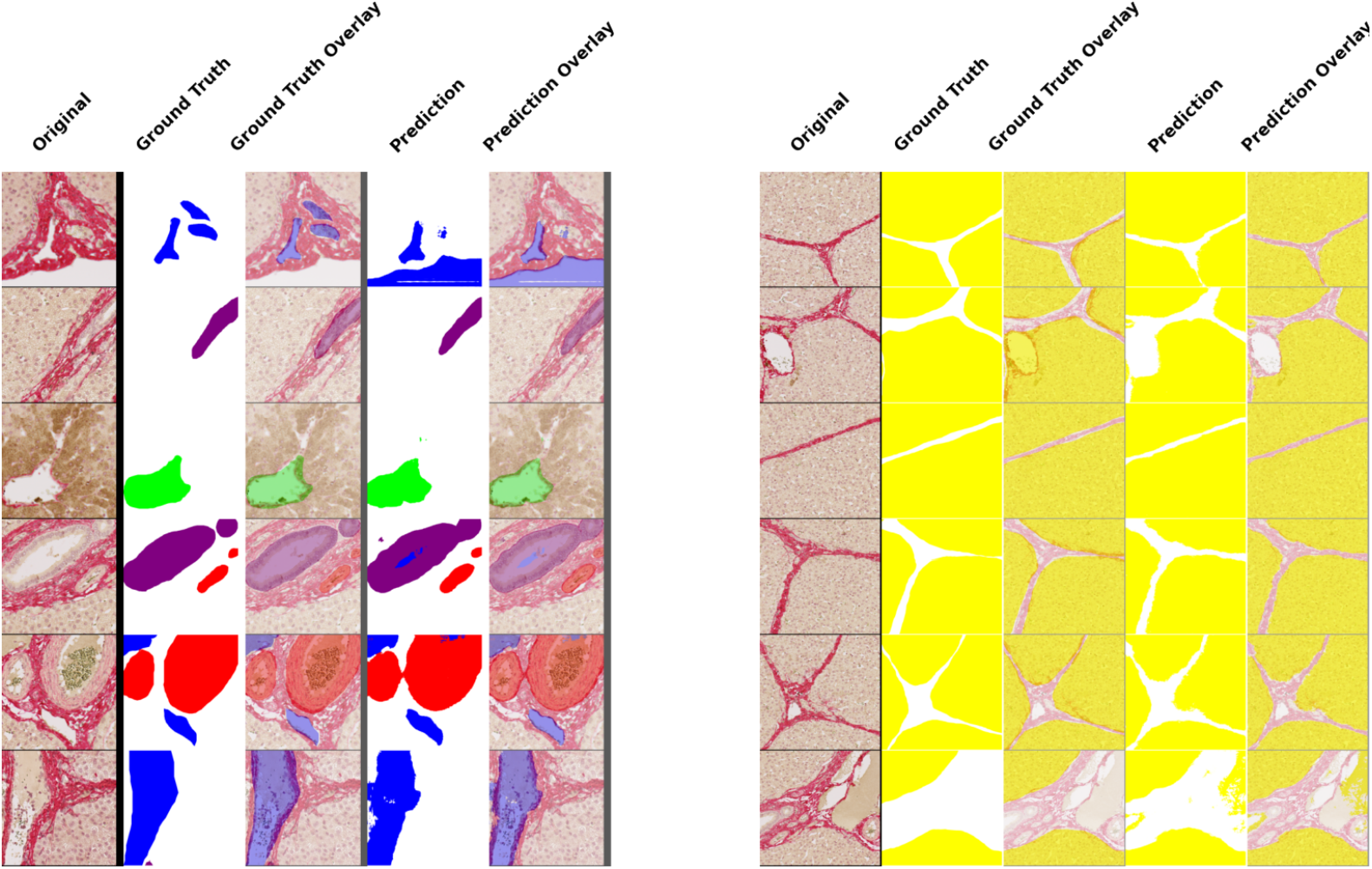
Visualization of automated segmentation of liver histology. Left panel: Blood vessel segmentation with ground truth and predictions. Right panel: Hepatic lobule segmentation with ground truth and predictions. Each row represents a different test sample. Columns show: (1) Original GS-PSR stained tissue, (2) Ground truth annotation, (3) Ground truth overlay, (4) Model prediction, (5) Prediction overlay

The confusion matrix in the **Figure 6** illustrates the performance of both models across all classes, emphasizing their classification accuracy and error patterns. Using the hepatic vessel model, the background is classified with very high accuracy, showing strong model performance in distinguishing the background from the targeted histological structures. Central Veins also have high accuracy (96.7%), with minimal confusion (3.3%) primarily with background. Portal Veins have a lower accuracy (85.7%), with notable confusion (12.0%) with arteries, indicating overlap or similarity in features. The arteries are well-identified (93.9%), though 5.7% are misclassified as background and a small portion as the portal vein. The bile ducts have the lowest accuracy (83.2%), with a substantial portion (13.9%) misclassified as Artery, suggesting these two structures are particularly challenging for the model to differentiate.

For the lobule model, the background achieves excellent classification accuracy (93.5%), though with some confusion (6.5%) with lobule structures. The lobule class demonstrates strong performance (96.7%) with minimal misclassification (3.3%) as background, indicating the model effectively distinguishes lobular architecture from surrounding tissue.

**Table 4** provides Average Symmetric Surface Distance (ASSD) in micrometers for the five hepatic structures. Portal veins showed the highest segmentation error (24.50 *µ*m average) with the greatest variability, while central veins achieved the lowest error (5.10 *µ*m) with the most consistent performance. The remaining structures demonstrated intermediate accuracy levels ranging from 6.07-10.95 *µ*m ASSD.

**Table 4:** Average Symmetric Surface Distance (ASSD) statistics for anatomical structures. Distances are reported in *µ*m, converted from pixels using the original slide resolution of 227 nm/pixel.

| Structure | Avg. ASSD | Std. Dev. | Median | Min | Max |
| --- | --- | --- | --- | --- | --- |
| Portal Vein | 24.50 $\mu\text{m}$ | 27.19 $\mu\text{m}$ | 15.94 $\mu\text{m}$ | 1.94 $\mu\text{m}$ | 125.10 $\mu\text{m}$ |
| Artery | 10.95 $\mu\text{m}$ | 12.04 $\mu\text{m}$ | 3.89 $\mu\text{m}$ | 0.52 $\mu\text{m}$ | 43.80 $\mu\text{m}$ |
| Central Vein | 5.10 $\mu\text{m}$ | 2.96 $\mu\text{m}$ | 4.23 $\mu\text{m}$ | 1.27 $\mu\text{m}$ | 19.12 $\mu\text{m}$ |
| Bile Duct | 8.46 $\mu\text{m}$ | 7.84 $\mu\text{m}$ | 7.90 $\mu\text{m}$ | 0.93 $\mu\text{m}$ | 30.45 $\mu\text{m}$ |
| Lobule | 6.07 $\mu\text{m}$ | 7.13 $\mu\text{m}$ | 3.78 $\mu\text{m}$ | 0.96 $\mu\text{m}$ | 45.49 $\mu\text{m}$ |

### 3.2 Robustness

To assess the robustness of the algorithm towards technical issues possibly occurring in routine histological preparations, we applied the model to four additional sections of varying technical quality, as shown in **Figure 8**. We assessed the results visually by comparing original images (**Figure 8**, left panel) with the corresponding predicted segmentation (**Figure 8**, right panel).

**Figure 8.**
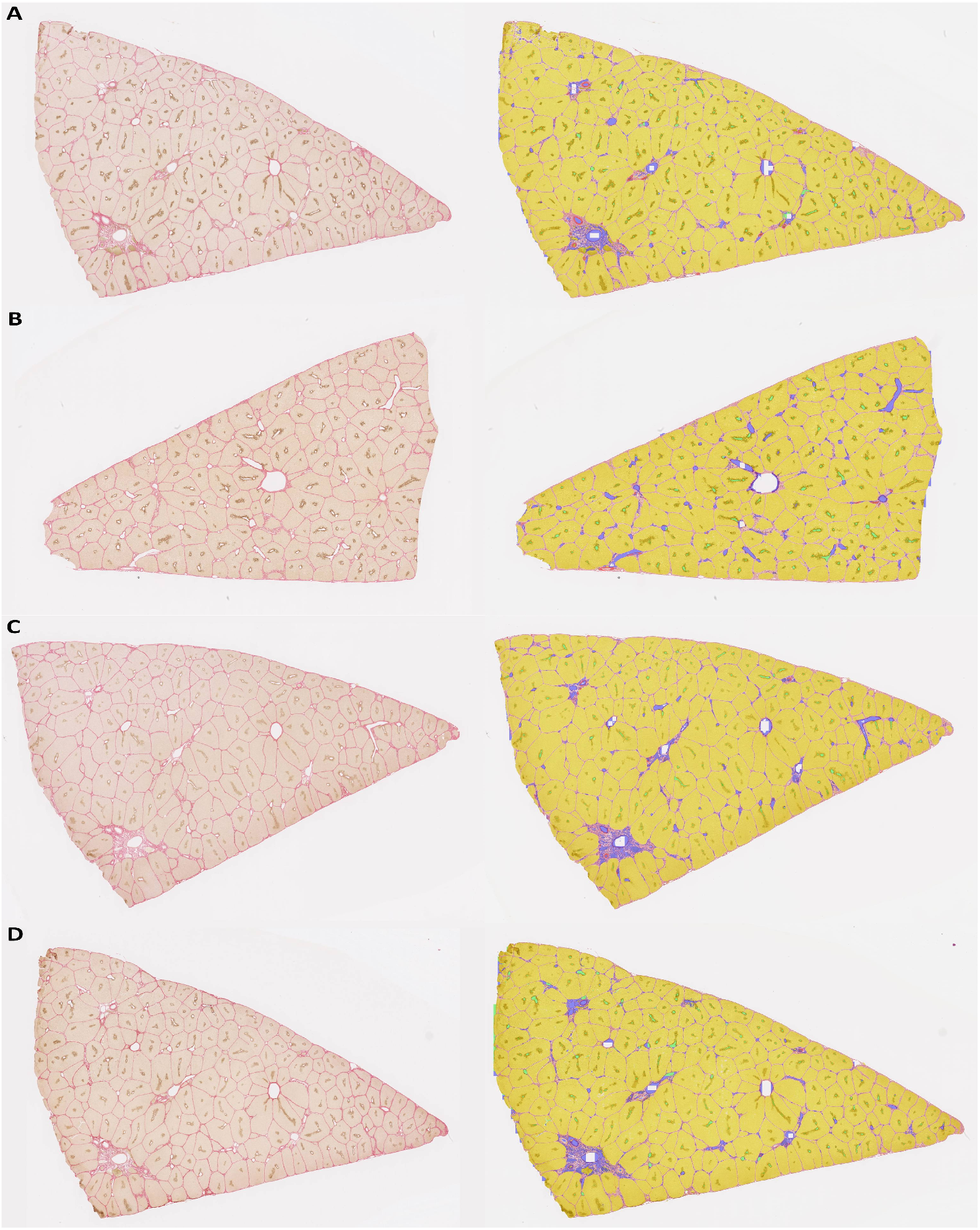
Automated segmentation results demonstrating algorithm robustness across varying tissue quality conditions. Four representative porcine liver tissue sections stained with GS-PSR are shown. Left panel: original histological images displaying varying degrees of technical quality. Right panel: corresponding automated segmentation results with color-coded tissue classification.

All the four sections’ images (**Figure 8A, Figure 8B, Figure 8C**, and **Figure 8D**) with high technical quality demonstrate optimal recognition of all targeted structures, yielding optimal segmentation results.

### 3.3 Morphometric analysis

Figure 9. illustrates the results for the morphometric analysis of the liver section depicted in **Figure 8C** with 1,473 hepatic structures classified as five distinct anatomical categories.

Portal veins comprised the largest group (n=492, 33.4%), followed by central vein (n=293, 19.9%), arteries (n=270, 18.3%), bile ducts (n=246, 16.7%), and lobules (n=172, 11.7%). Shape analysis showed lobules being more circular compared to e.g. portal veins as indicated by the higher respectively lower aspect ratio with a median of 0.65 compared to 0.45. Size measurements showed that lobules were substantially larger (median major feret diameter *≈* 1,000 *µ*m; area *≈* 450,000 *µ*m^2^) than vascular structures, with portal veins (median major feret diameter*≈* 105 *µ*m) exceeding arteries, bile ducts, and central veins (30–60 *µ*m range). These quantitative findings illustrate the structural organization of hepatic tissue, spanning multiple scales from large lobules that define tissue architecture to fine vascular structures for microcirculation.

**Figure 9.**
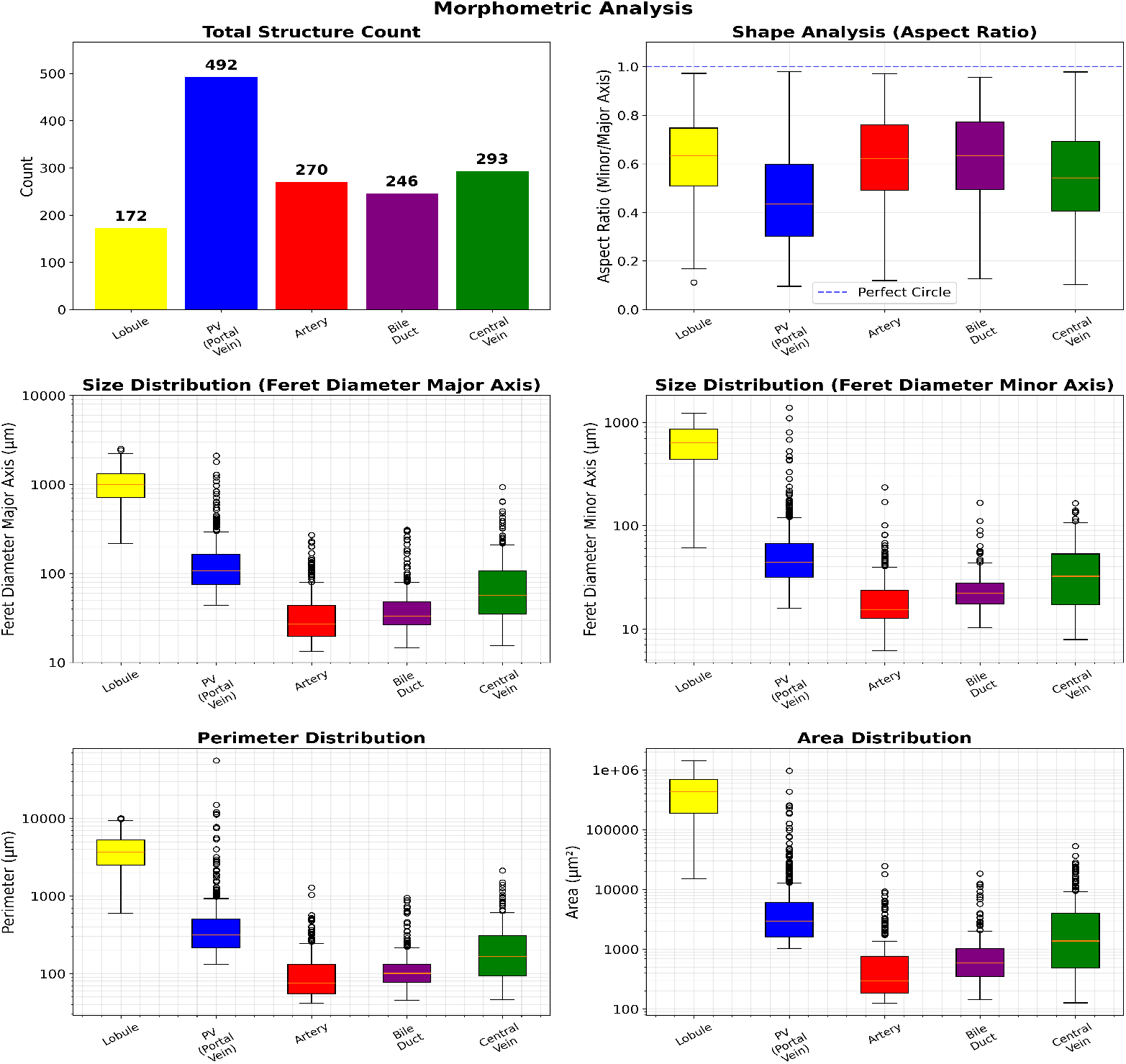
Quantitative parameters derived from histological image analysis include total structure count, shape aspect ratio (minor/major axis), feret diameters (major and minor), perimeter, and area. Aspect ratio analysis indicates deviation from circularity, especially in vessels and bile ducts. Logarithmic scaling was applied to perimeter and area plots for clarity. In the box plots, boxes represent quartiles Q1 and Q3. The upper whisker and lower whisker extend to the last data point less than Q3 + 1.5*·*IQR and the first data point greater than Q1 *−* 1.5*·*IQR, respectively. IQR denotes interquartile range (Q3–Q1).

## 4 Discussion

Comprehensive morphometric analysis of hepatic morphology calls for the identification and quantification of all histological structures. This study presents a deep-learning framework for segmenting key liver structures, including portal veins, hepatic arteries, bile ducts, central veins, and lobules, in whole slide scans of porcine liver tissue. Our approach did achieve strong segmentation performance independent of size across different types of histological structures within the liver.

### 4.1 Technical Comparison with Existing Approaches

Current hepatic tissue segmentation approaches predominantly target isolated anatomical components, such as bile ducts or portal tracts, limiting their applicability in comprehensive liver tissue analysis (see **Table 5**).

**Table 5:** Technical comparison of existing approaches for vessel segmentation.

| Study | Approach | Origin | Tissue Condition | Staining | Detected Structures | Performance Metrics |
| --- | --- | --- | --- | --- | --- | --- |
| (author?)<br>[SCL<sup>+</sup>22] | Multi-Scale Attention CNN | Human liver biopsies | Fibrotic (Chronic Hepatitis) | Masson Stained | Bile Duct | DSC: 0.84<br>IoU: 0.73<br>TPR: 0.86 |
| (author?) [YSJ<sup>+</sup>22] | MUSA-UNet | Human transplantation biopsy | Transplant biopsy | HE, Masson Trichrome | Portal Field | DSC: 0.89<br>IoU: 0.79<br>TPR: 0.85 |
| (author?)<br>[BLW<sup>+</sup>22] | Cascade-R-CNN | Mouse liver | Healthy, Resected, Steatotic | HE, GS | Lobule, Portal Field, Central Vein | DSC: 0.79–0.81<br>IoU: 0.66–0.68<br>TPR: 0.81–0.84 |
| (author?)<br>[RWL<sup>+</sup>23] | MobileNet-v2 | Mouse liver | Healthy | GS and zoned markers | Central Vein, Portal Vein | DSC: 0.81–0.89<br>IoU: 0.89–0.94 |
| Bafna et al. (2025) | Weight boosting UNet | Pig liver | Healthy | GS-PSR | Portal Vein, Central Vein, Artery, Bile Duct, Lobule | DSC: 0.75–0.96<br>IoU: 0.63–0.93<br>TPR: 0.71–0.94 |

Four different strategies using deep learning networks for vessel segmentation in WSIs of liver histology were recently reported: Multi-scale attention convolutional network built on a ResNet-101 backbone with the atrous spatial pyramid pooling (ASPP) [SCL^+^22]. This technology builds on dual-magnification input processing with attention mechanisms to enhance feature maps from high-magnification images using low-magnification contextual information. The advantage is the integration of multi-scale spatial information via decoder architecture. However, high computational resource requirements and limited generalization to other vascular and lobular structures represent limitations. Similar to the attention mechanism-based approach in [SCL^+^22], [YSJ^+^22] proposed the Multiple Up-sampling and Spatial Attention guided U-Net model (MUSA-UNet), which the specifically adapted for portal tract region identification and quantification in transplant biopsy whole-slide images. This modified U-Net architecture incorporates multi-scale attention modules to effectively capture the complex morphological variations present in transplant biopsies. This strategy enhances feature representation and segmentation accuracy, but requires high computational resources due to the integration of attention mechanisms.

The other two approaches are rather different from these encoder-decoder strategies. [BLW^+^22] employed Cascade R-CNN with ResNeXt101 backbone for simultaneous detection of portal fields and central veins using a tile-based approach. Unlike attention-based segmentation networks that enhance feature representation via spatial weighting across scales, the Cascade R-CNN performs structured, multi-stage object detection by the bounding box approach without relying on learned attention maps. [RWL^+^23] utilized MobileNet-v2 for lightweight vessel detection in healthy tissue. Both approaches rely on object detection paradigms rather than semantic segmentation frameworks. In addition, the approach differs from [BLW^+^22] as it employs tissue positioning system (TPS), a lightweight semantic segmentation paradigm to generate continuous zonal masks focusing on positional mapping rather than discrete detection.

For lobular segmentation, methodological approaches have evolved from basic geometric principles to sophisticated multi-modal integration (see **Table 6**). [SHS^+^16] utilised a distance-based approach, employing computational geometry to define lobular boundaries through proximity measurements to vascular structures in steatotic mouse liver tissue. Building upon this framework, [LBA^+^24] introduced distance-based joint zonated quantification. This method combines the geometric principles of distance-based approaches with multi-stain integration capabilities, achieving enhanced zonation mapping accuracy. However, it comes at the cost of substantially increased computational complexity compared to single-parameter approaches.

**Table 6:**
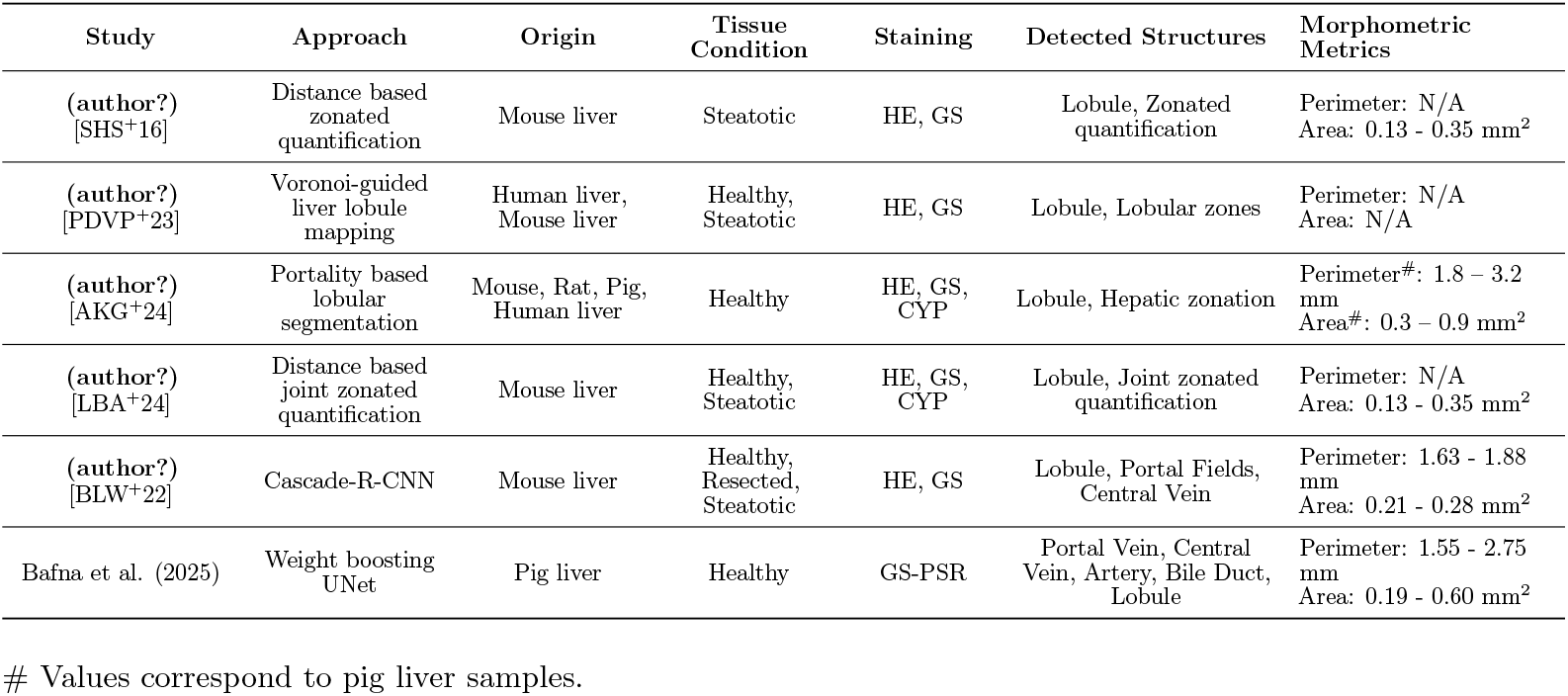
Technical comparison of existing approaches for lobular segmentation.

| Study | Approach | Origin | Tissue Condition | Staining | Detected Structures | Morphometric Metrics |
| --- | --- | --- | --- | --- | --- | --- |
| (author?)<br>[SHS<sup>+</sup>16] | Distance based zoned quantification | Mouse liver | Steatotic | HE, GS | Lobule, Zoned quantification | Perimeter: N/A<br>Area: 0.13 - 0.35 mm<sup>2</sup> |
| (author?)<br>[PDVP<sup>+</sup>23] | Voronoi-guided liver lobule mapping | Human liver, Mouse liver | Healthy, Steatotic | HE, GS | Lobule, Lobular zones | Perimeter: N/A<br>Area: N/A |
| (author?)<br>[AKG<sup>+</sup>24] | Portality based lobular segmentation | Mouse, Rat, Pig, Human liver | Healthy | HE, GS, CYP | Lobule, Hepatic zonation | Perimeter#: 1.8 – 3.2 mm<br>Area#: 0.3 – 0.9 mm<sup>2</sup> |
| (author?)<br>[LBA<sup>+</sup>24] | Distance based joint zoned quantification | Mouse liver | Healthy, Steatotic | HE, GS, CYP | Lobule, Joint zoned quantification | Perimeter: N/A<br>Area: 0.13 - 0.35 mm<sup>2</sup> |
| (author?)<br>[BLW<sup>+</sup>22] | Cascade-R-CNN | Mouse liver | Healthy, Resected, Steatotic | HE, GS | Lobule, Portal Fields, Central Vein | Perimeter: 1.63 - 1.88 mm<br>Area: 0.21 - 0.28 mm<sup>2</sup> |
| Bafna et al. (2025) | Weight boosting UNet | Pig liver | Healthy | GS-PSR | Portal Vein, Central Vein, Artery, Bile Duct, Lobule | Perimeter: 1.55 - 2.75 mm<br>Area: 0.19 - 0.60 mm<sup>2</sup> |

[PDVP^+^23] advanced the methodology by implementing Voronoi-guided mapping strategies. These improved upon simple distance calculations by using tessellation algorithms to create more physiologically accurate lobular zone partitions across both human and mouse samples under healthy and steatotic conditions.

More recent developments have shifted toward comprehensive multi-species applicability. Portality-based segmentation extends beyond the species-limited approaches of earlier methods to encompass mouse, rat, pig, and human tissues [AKG^+^24]. This approach further differentiates itself by integrating multiple staining protocols (HE, GS, and CYP) for enhanced lobular analysis. In contrast, previous methods relied predominantly on single-stain approaches. [BLW^+^22] estimated lobular dimensions by utilizing the identified positions of the PF and CV, as mentioned in **Table 5** describing their vessel segmentation framework.

In contrast, the present work proposes an integrated multi-class vessel segmentation framework paired with lobule segmentation. The methodology utilizes GS and PSR staining on porcine liver sections to improve the contrast of key anatomical features, such as collagen-dense vessel walls and lobular boundaries. In contrast to other studies mainly focusing on one or few selected structures, this framework simultaneously segments five hepatic components—lobules, portal veins, central veins, hepatic arteries, and bile ducts—within a single pipeline. The weight-boosting mechanism specifically targets class imbalance challenges inherent in the hepatic vessel detection with metrics comparable to single-structure methods while maintaining multi-target detection capabilities. Our comprehensive approach represents a substantial advancement in automated hepatic histological analysis, unified for complex multi-structure segmentation tasks. The proposed weight boosting U-Net mechanism demonstrates substantial advancements in terms of vessel segmentation, but also has several biological and technical constraints limiting its current applicability and generalizability. The model was developed using WSI of GS-PSR stained sections from healthy porcine livers which represents a narrow scope of application. Four critical parameters define the operational boundaries of our segmentation framework: species specificity, staining protocol dependency, tissue condition requirements, and section quality standards. Each parameter presents unique challenges that must be addressed to achieve strong performance across diverse scenarios.

### 4.2 Future directions

Taken together, our pipeline enables quantitative vascular architecture assessment and serves as a foundation extending the applicability of the algorithm to other conditions. As a next step, we aim to perform quantitative morphometric analysis across species (mouse, rat, human) and across different stainings, focusing on both tissue morphology and marker protein expression. Further plans include the extension to the total quantification of liver pathologies (hepatic steatosis, fibrosis/cirrhosis, necrosis, inflammation, regeneration), but also the inclusion of zonated quantification of these morphologic changes and, respectively, zonated expression of marker proteins.

To achieve this goal, the key prerequisite on the biological side is the generation of comprehensive annotated datasets across different species, stainings, technical quality, and liver pathologies. The following four challenges must be addressed. The generalization to multiple species requires enhancing lobular structure identification in species lacking lobular septa. The incorporation of diverse staining methods calls for the implementation of stain normalization, followed by training on differentially stained sections. The inclusion of different tissue conditions requires the annotation of comprehensive datasets representing an array of pathological states. The applicability to images from sections of suboptimal quality depends on the design of preprocessing pipelines and architectures resilient to artifacts and sectioning flaws.

On the image analysis side, the key steps include staining normalization techniques and domain adaptation methods. Stain normalization refers to the computational adjustment of histological images to reduce color variability introduced by differences in staining protocols or scanners, ensuring consistent feature representation across samples. In this context, domain adaptation addresses cross-species variability by enabling models trained on one species’ tissue morphology to generalize effectively to others, such as transferring between human and porcine liver samples.

## 5 Conclusion

Automated segmentation of hepatic vessels and lobular structures in whole-slide histological images is a fundamental task in computational liver histology, enabling detailed anatomical analysis and supporting downstream image-based quantification. The primary challenge is to correctly identify complex vascular structures (portal veins, central veins, arteries, bile ducts, and lobules) with varying diameters throughout the liver parenchyma.

We implemented two U-Net variants enhanced with a weight-boosting mechanism to segment and classify specific hepatic vessel types in whole-slide porcine liver images. Our model is achieving comparable accuracy in specific vessel type classification as reported for general vessel segmentation, while allowing structure-wise morphometric analysis. Nevertheless, the applicability is currently restricted to WSI from GS-PSR-stained high-quality sections from healthy porcine livers.

A coordinated effort to address this main limitation and to extend applicability would substantially advance this field, potentially leading to a robust approach that adapts and transfers efficiently across species, stainings, and pathologies.

## Supporting information

Artery_8C

Artery_8D

BileDuct_8C

BileDuct_8D

CentralVein_8C

CentralVein_8D

Lobule_8C

Lobule_8D

Morphometry_D

PortalVein_8C

PortalVein_8D

## Ethics Statement

The work with animals was conducted under the law of the Czech Republic, which is in line with the legislation of the European Union. The procedure protocol was approved by the Ministry of Education, Youth and Sport of the Czech Republic (approval no. MSMT-15629/2020-4)..

## Author Contributions

MB contributed to the conceptualization, software development, data curation, formal analysis, investigation, methodology, validation, visualization, original draft preparation, and review and editing of the manuscript. MA contributed to data curation, investigation, methodology, validation, visualization, and manuscript review and editing. VL provided resources. VM contributed resources and participated in manuscript review and editing. SS was involved in conceptualization, supervision, project administration, manuscript review and editing, and funding acquisition. MK contributed to methodology, manuscript review and editing, and funding acquisition. UD contributed to conceptualization, data curation, formal analysis, investigation, methodology, validation, visualization, supervision, project administration, resource provision, and manuscript review and editing.

## Supplemental Data

Supplementary materials related to **Figure 8** include:

- Five **CSV** files containing spatial and morphometric data for each segmented structure (artery, bile duct, central vein, portal vein, and lobule), corresponding to panels C and D.
- An **image summarizing morphometric analyses** of area, perimeter, and Feret diameter for all segmented structures in panel D.

These materials provide additional quantitative and visual context supporting the segmentation results presented in Figure 8.

## Data Availability Statement

The datasets generated and analyzed for this study can be found in the Zenodo repository: https://doi.org/10.5281/zenodo.17041227.

## References

[ADEK19] S. K. Asrani, H. Devarbhavi, J. Eaton, and P. S. Kamath. Burden of liver diseases in the world. The Lancet Gastroenterology & Hepatology, 4(3):175–191, 2019.

[AKG+24] Mohamed Albadry, Jonas Küttner, Jan Grzegorzewski, Olaf Dirsch, Eva Kindler, Robert Klopfleisch, Vaclav Liska, Vladimira Moulisova, Sandra Nickel, Richard Palek, Jachym Rosendorf, Sylvia Saalfeld, Utz Settmacher, Hans-Michael Tautenhahn, Matthias König, and Uta Dahmen. Cross-species variability in lobular geometry and cytochrome p450 hepatic zonation: insights into cyp1a2, cyp2d6, cyp2e1 and cyp3a4. Frontiers in Pharmacology, Volume 15 - 2024, 2024.

[BLW+22] Daniel Budelmann, Hendrik Laue, Nick Weiss, Uta Dahmen, Lorenza A. D’Alessandro, Ina Biermayer, Ursula Klingmüller, Ahmed Ghallab, Reham Hassan, Brigitte Begher-Tibbe, Jan G. Hengstler, and Lars Ole Schwen. Automated detection of portal fields and central veins in whole-slide images of liver tissue. Journal of Pathology Informatics, 13:100001, 2022.

[CK21] M. Ciecholewski and M. Kassjański. Computational methods for liver vessel segmentation in medical imaging: A review. Sensors, 21(6):2027, 2021.

[CS16] John M. Cullen and Margaret J. Stalker. Chapter 2 - liver and biliary system. In M. Grant Maxie, editor, Jubb, Kennedy & Palmer’s Pathology of Domestic Animals: Volume 2 (Sixth Edition), pages 258–352.e1. W.B. Saunders, sixth edition edition, 2016.

[GGH+22] A. Goode, B. Gilbert, J. Harkes, D. Jukic, and M. Satyanarayanan. Openslide: A vendor-neutral software foundation for digital pathology. Journal of Pathology Informatics, 3(1):1–9, 2022.

[HAT+23] Georg Hille, Shubham Agrawal, Pavan Tummala, Christian Wybranski, Maciej Pech, Alexey Surov, and Sylvia Saalfeld. Joint liver and hepatic lesion segmentation in mri using a hybrid cnn with transformer layers. Computer Methods and Programs in Biomedicine, 240:107647, 2023.

[HJH+24] Georg Hille, Tameem Jahangir, Janine Hürtgen, Rober Kreher, and Sylvia Saalfeld. Deep learning-based liver vessel segmentation. Current Directions in Biomedical Engineering, 10(1):29–32, 2024.

[Inv13] Pietro Invernizzi. Liver auto-immunology: The paradox of autoimmunity in a tolerogenic organ. Journal of Autoimmunity, 46:1–6, 2013. Liver Autoimmunity.

[IPK+18] Fabian Isensee, Jens Petersen, Andre Klein, David Zimmerer, Paul F. Jaeger, Simon Kohl, Jakob Wasserthal, Gregor Koehler, Tobias Norajitra, Sebastian Wirkert, and Klaus H. Maier-Hein. nnu-net: Self-adapting framework for u-net-based medical image segmentation, 2018.

[JBB79] L. C. Junqueira, G. Bignolas, and R. R. Brentani. Picrosirius staining plus polarization microscopy, a specific method for collagen detection in tissue sections. The Histochemical Journal, 11(4):447–455, 1979.

[LBA+24] Hendrik Oliver Arp Laue, Daniel Budelmann, Mohamed Albadry, Christiane Engel, Nick Weiss, Uta Dahmen, and Lars Ole Schwen. Joint zonated quantification of multiple parameters in hepatic lobules. 2024.

[LYL+14] Raed Lattouf, Ronald Younes, Didier Lutomski, Nada Naaman, Gaston Godeau, Karim Senni, and Sylvie Changotade. Picrosirius red staining: a useful tool to appraise collagen networks in normal and pathological tissues. Journal of Histochemistry & Cytochemistry, 62(10):751–758, 2014.

[MSN+12] F. Meurens, A. Summerfield, H. Nauwynck, L. Saif, and V. Gerdts. The pig: a model for infectious diseases research. Trends in Microbiology, 20(1):50–57, 2012.

[Nor79] MD Norenberg. Distribution of glutamine synthetase in the rat central nervous system. Journal of Histochemistry & Cytochemistry, 27(3):756–762, 1979. PMID: 39099.

[PDVP+23] Cédric Peleman, Winnok H De Vos, Isabel Pintelon, Ann Driessen, Annelies Van Eyck, Christophe Van Steenkiste, Luisa Vonghia, Joris De Man, Benedicte Y De Winter, Tom Vanden Berghe, et al. Zonated quantification of immunohistochemistry in normal and steatotic livers. Virchows Archiv, 482(6):1035–1045, 2023.

[RWL+23] Ruichen Rong, Yonglong Wei, Lin Li, Tao Wang, Hao Zhu, Guanghua Xiao, and Yunguan Wang. Image-based quantification of histological features as a function of spatial location using the tissue positioning system. EBioMedicine, 94, 2023.

[SCL+22] Chun-Han Su, Pau-Choo Chung, Sheng-Fung Lin, Hung-Wen Tsai, Tsung-Lung Yang, and Yu-Chieh Su. Multi-scale attention convolutional network for masson stained bile duct segmentation from liver pathology images. Sensors, 22(7), 2022.

[SHS+16] Lars Ole Schwen, André Homeyer Michael Schwier, Uta Dahmen, Olaf Dirsch, Arne Schenk, Lars Kuepfer, Tobias Preusser, and Andrea Schenk. Zonated quantification of steatosis in an entire mouse liver. Computers in biology and medicine, 73:108–118, 2016.

[TGW17] Elijah Trefts, Maureen Gannon, and David H. Wasserman. The liver. Current Biology, 27(21):R1147–R1151, 2017.

[YSJ+22] Hanyi Yu, Nima Sharifai, Kun Jiang, Fusheng Wang, George Teodoro, Alton B. Farris, and Jun Kong. Artificial intelligence based liver portal tract region identification and quantification with transplant biopsy whole-slide images. Computers in Biology and Medicine, 150:106089, 2022.

