## Supplementary figures and images for "Automated Segmentation of Hepatic Vessels and Lobules in Whole-Slide Images Using U-Net Models"

### Morphometry_D

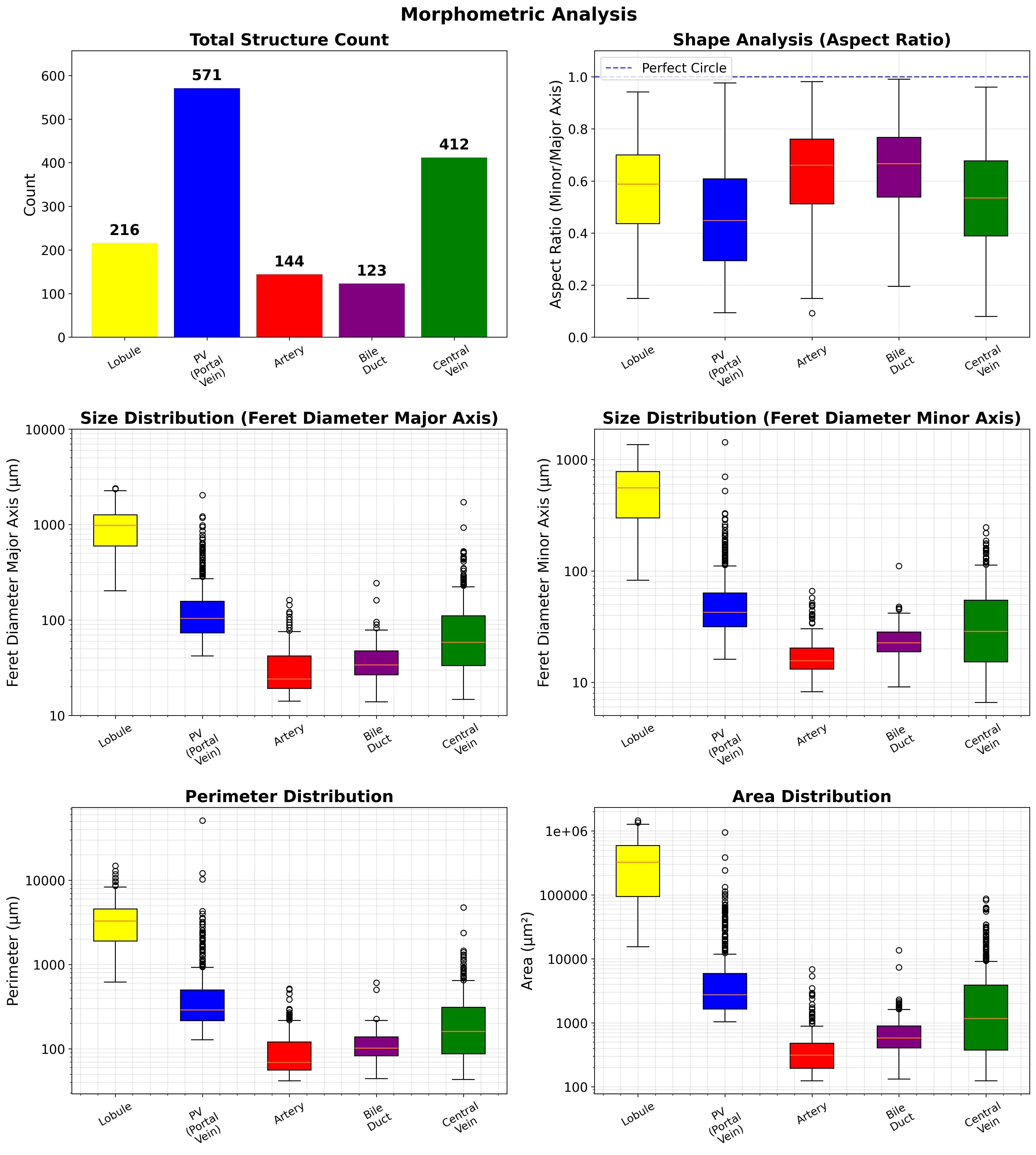
